# RIBEX: Predicting and Explaining RNA Binding Across Structured and Intrinsically Disordered Regions (IDR)-rich Proteins

**DOI:** 10.64898/2026.03.13.711639

**Authors:** Samuele Firmani, Felix Steinbauer, Gjergji Kasneci, Annalisa Marsico, Marc Horlacher

## Abstract

**Motivation:** RNA-binding proteins (RBPs) regulate post-transcriptional processes, yet many remain undiscovered because RNA-binding activity often occurs outside canonical RNA-binding domains (RBDs), including within intrinsically disordered regions (IDRs) or through protein complexes. Computational methods can help identify novel RBPs, but approaches relying solely on sequence-derived features or ignoring the cellular interaction context are limited in capturing the complexity of RNA-binding behavior. To date, no framework rigorously integrates both sequence information and protein interaction context for RBP prediction.

**Results:** We introduce RIBEX, a multimodal framework that combines protein language model (pLM) embeddings with protein interactome topology to improve RBP prediction and interpretation. Specifically, we integrate sequence representations with graph-derived positional encodings (PE) from the human STRING protein–protein interaction (PPI) network. PE are computed using Personalized PageRank, reduced with principal component analysis, and fused with pooled sequence embeddings through FiLM conditioning, while Low-Rank Adaptation (LoRA) enables parameter-efficient task adaptation. Across both an annotation-based benchmark and experimental RNA Interactome Capture (RIC) dataset, PE consistently improves predictive performance, indicating that interactome topology provides complementary information beyond sequence features. LoRA adaptation of ESM2-650M further yields larger gains than simply scaling frozen backbone size. RIBEX outperforms state-of-the-art methods such as RBP-TSTL and HydRA, particularly on challenging subsets including proteins lacking canonical RBDs and those enriched in IDRs. For interpretability, we combine sequence-level computational alanine scanning with network-level positional-encoding ablation and inverse-PCA mapping, recovering known RNA-binding domains, IDR-associated contributions, and functional interactome communities linked to RBP predictions.

## Introduction

The regulation of RNA transcripts is orchestrated by a diverse class of molecules known as RNA-Binding Proteins (RBPs). Traditionally, RBPs were defined by canonical RNA-Binding Domains (RBDs), however, high-throughput methods like RNA Interactome Capture (RIC) uncovered a large “hidden” layer of the interactome (6), revealing hundreds of previously unrecognized and non-canonical RBPs. Many of these lack classical RBDs and instead rely on low-complexity, Intrinsically Disordered Regions (IDRs), now recognized as key mediators of RBP–RNA interactions (10).

Although RBPs have been systematically annotated across multiple species in diverse databases (2), their repertoire remains far from complete. RIC enables RBP identification on a large scale, but it provides only a condition-specific snapshot RNA–protein interactions, often missing low-abundance RBPs, proteins that bind non-polyadenylated RNAs, and RBPs expressed in other cell types or experimental contexts (e.g., a RIC screen in HEK293 cells will not detect RBPs restricted to neurons or liver). Computational methods that learn the sequence and structural “grammar” of binding can bridge these gaps by providing a condition-independent, genome-wide scan of the proteome and predicting RBPs where experimental data are unavailable. However, predicting novel RBPs remains a significant challenge, as many RBPs lack the evolutionary conservation or structural signatures relied upon by classical computational methods.

Early computational RBP prediction methods either trained supervised classifiers on sequence and structural features (e.g., biochemical properties, evolutionary conservation) or relied on known RBDs and motifs (16; 21).

However, domain-based approaches are limited, as about one third of experimentally identified RBPs lack recognizable RNA-binding homology. Tools such as RBPPred (26) used Support Vector Machines (SVMs) with Position-Specific Scoring Matrices (PSSMs) to incorporate evolutionary information, but at high computational cost. This motivated Deep-RBPPred, which leveraged Convolutional Neural Networks (CNNs) to learn local biophysical patterns more efficiently (27). TriPepSVM advanced the field by moving from global homology and complex structural features to modeling short amino acid motifs—the local grammar of binding—specifically tri-peptides, and was particularly effective at identifying RBPs with disordered regions, as basic- and aromatic-rich tri-peptides are enriched in IDRs of known binders (5). However, purely sequence-based approaches remain limited when long-range interactions are essential for protein function.

A large class of pretrained Protein Language Models (pLMs), such as ESM-2/3 (15), treat amino acid sequences like natural language and are trained self-supervised on large protein sequence datasets. They capture contextual, local, and global sequence features, as well as latent structural and functional information, making them the leading framework for transfer learning in protein property and structure prediction (comprehensive overview in (25)). A landmark study, RNA-Binding Protein Two-Stage Transfer Learning (RBP-TSTL) (18), introduced a two-stage transfer learning approach to improve genome-scale RBP prediction. It leverages high-dimensional embeddings from pLMs with limited task-specific data: a pre-trained ESM model first extracts generic sequence features, which are then fine-tuned on annotated RBPs to capture RNA-binding-specific signatures.

However, even pLM-based models face a limitation: they primarily learn from the internal “context” of a single protein sequence and often tend to overlook the extrinsic context—the protein’s cellular environment. RBPs tend to cluster in functional “neighborhoods” within the interactome (e.g., spliceosome, ribosome), and a protein’s propensity to bind RNA is often reflected in its Protein–Protein Interaction (PPI) network. Tools such as Support Vector Machine obtained from neighborhood associated RBPs (SONAR)(4) were among the first to demonstrate that analyzing PPI association patterns could discover novel RBPs that sequence-only methods missed. More recently, Hybrid Ensemble Classifier for RNA-Binding Proteins (HydRA)(12) integrated this interaction context with deep learning, proving that the “neighborhood” of a protein provides a powerful signal for its RNA-binding capacity. Despite these advancements, a gap remains for non-canonical RBPs, and current models often fail to capture the synergy between a protein’s global network position and its local, disordered binding motifs.

Recent work has shown that well-designed Positional Encodings (PEs) capturing a node’s structural position in a graph can boost prediction performance (8), a principle not yet widely applied to PPI networks in genomics. For RBP predictions, Positional Encodings (PEs) are particularly valuable for non-canonical RBPs lacking structured RNA-binding domains: by encoding a protein’s topological role (e.g., hub or peripheral bridge), they can distinguish intrinsically disordered proteins near known RBPs—which are more likely functional binders—from proteins with similar sequences but different interactome roles.

In this work, we present RIBEX (RIBEX), a simple yet powerful multimodal framework that integrates protein language modeling with network context to improve RNA-binding prediction. By combining parameter-efficient Low-Rank Adaptation (LoRA) fine-tuning of ESM-2 with positional encoding from the PPI network, we merge intrinsic sequence features with the external context of the interactome. This approach preserves the broad biological knowledge encoded in foundation models while enhancing the detection of non-canonical and IDR-mediated RBPs, achieving superior performance compared with state-of-the-art methods such as RBP-TSTL and HydRA. We further interpret predictions using *in silico* alanine scanning (13) to identify key sequence regions—such as IDRs or specific domains—driving RNA-binding predictions. These signals align with structural confidence patterns from AlphaFold, supporting the biological plausibility of our approach. Beyond predictive accuracy and interpretability, our work clarifies when task-specific adaptation is beneficial and highlights the complementary value of network information in uncovering functionally relevant RBPs.

## Materials and Methods

We formulate RBP identification as a binary classification task over human proteins, using (i) the amino-acid sequence of each protein and (ii) its topological context in a human PPI graph derived from STRING (24).

### Data Collection and Pre-Processing

Two primary datasets are used for model training and further analysis (i) the annotation-based set from (5), and (ii) an experimental RIC dataset from RBPbase (10). To align with our target domain, both are filtered to the human proteome (taxon ID 9606). In the RIC dataset, we classify proteins as RBP targets (positive class) if verified in at least one (otherwise negative) high-throughput assay from well-characterized human cell lines (HuH7, HeLa, JURKAT, MCF7, and HEK293). We further integrate functional annotations for RNA-binding (QuickGO:0003723) and IDRs from InterPro (3) and MobiDB (20), respectively. Following retrieval, protein sequences—capped at 1,024 residues, as described in (18)—are transformed into dense embeddings using ESM-2 (650M and 3B variants) (15) and ProtT5-XL (3B) (9). In addition, we benchmark against the dataset from HydRA (12), which combines protein sequence information and protein interaction context features. Positive examples were collected from experimentally supported RBP annotations and large-scale RNA interactome studies, while negative examples correspond to proteins without RNA-binding evidence. The dataset also integrates PPI and functional association networks to capture the cellular context of proteins. For more details on dataset composition and split statistics, see Suppl. Section A.1 and Section A.2.

To avoid inflated performance due to homology leakage, we perform all **dataset partitioning** at the *homology-cluster* level rather than at the individual-sequence level. We first cluster the full sequence set with MMseqs2 (23) using a minimum sequence identity of 20% and bidirectional coverage ≥ 50%. Each protein is assigned the identifier of its cluster representative (centroid), and clusters are treated as indivisible units during splitting (singletons are treated as clusters of size one), guaranteeing that no homologous proteins appear on both sides of a split. For the RIC and (5) datasets, only proteins for which embeddings are available across all compared pLMs are retained, forming a common gene pool shared by all models to ensure model-agnostic and directly comparable evaluation. From this shared pool, 10% of the data is held out; this hold-out set is further partitioned into a validation fold (one third) used for early stopping and epoch selection, and a final test fold (two thirds) used for performance evaluation. Dataset composition and split sizes for both benchmarks are summarised in Tables 1 and 2. For the HydRA benchmark, the official held-out test set published in the HydRA repository (12) is used directly as the evaluation set, while the remaining proteins serve as training data. Per-seed validation proteins for epoch selection are obtained by bootstrap resampling with replacement from the test set, treating the out-of-bag proteins as the validation fold for that seed. Split statistics for the HydRA benchmark are reported in Suppl. Table 3.

### Model Architecture

RIBEX integrates sequence-based and network-derived node positions to predict human RNA-binding proteins. A pre-trained pLM first encodes the primary sequence into high-dimensional, contextualized residue embeddings. These features are subsequently condensed via masked mean pooling to generate a fixed-length protein representation. To ensure computational efficiency, we employ LoRA (11) for model fine-tuning. While the core pLM weights remain frozen, trainable low-rank matrices are integrated into the attention layers and optimized for the classification task (Figure 1). Concurrently, task-specific layers for feature fusion (Section 2.3) and final prediction are trained in an end-to-end fashion.

**Figure 1.**
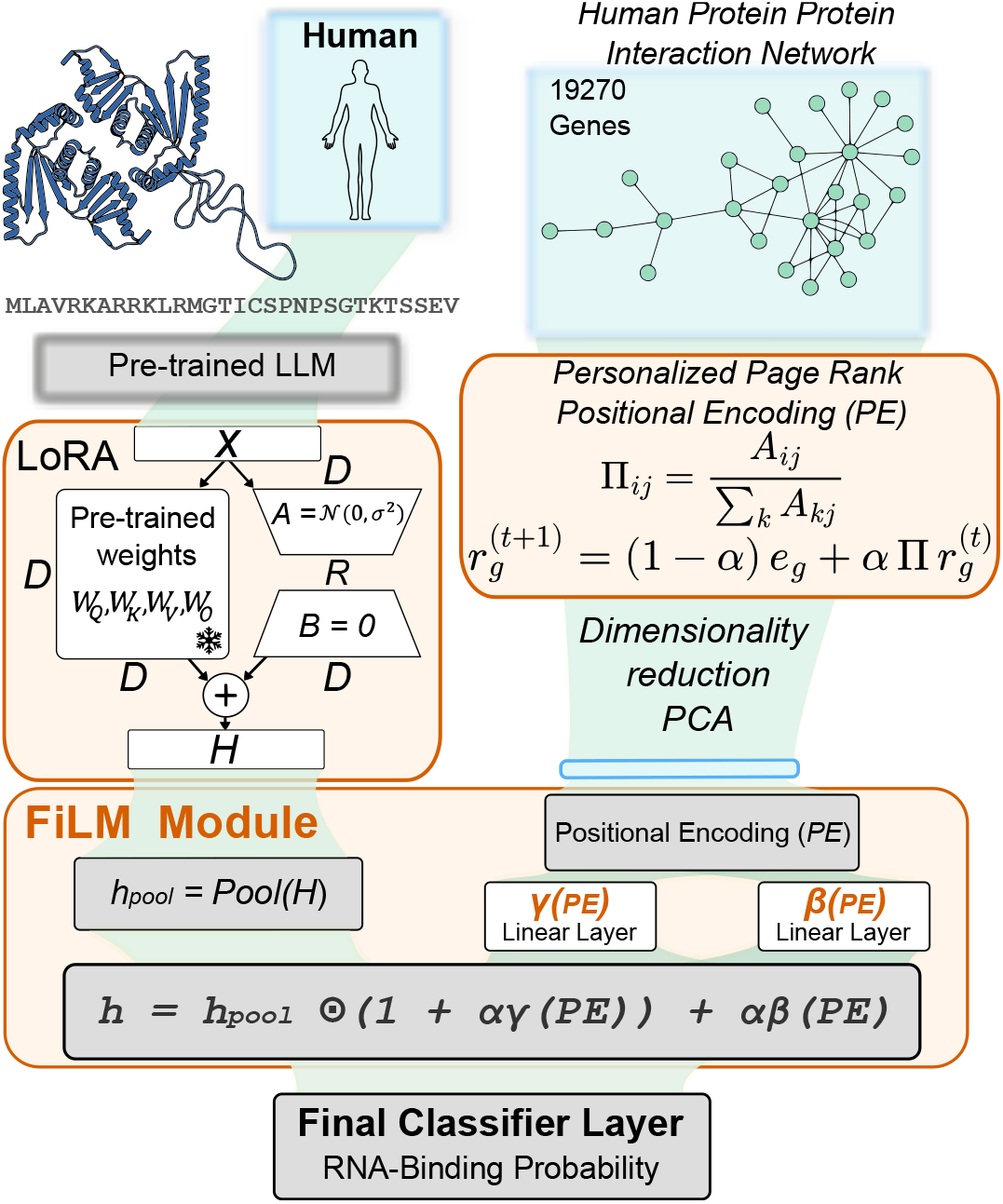
Overview of RIBEX. A protein language model encodes the amino-acid sequence, while graph-derived positional encodings from the human STRING PPI network (Personalized PageRank (PPR) followed by Principal Component Analysis (PCA)) capture interactome context. A FiLM layer conditions the pooled sequence representation on the positional encoding, and a classifier predicts RNA-binding probability.

### Positional Encoding and Feature Integration

Beyond sequence data, we incorporate the broader network context of each protein through its position in the human PPI graph. We characterize the local topology of each node using PPR vectors (17). Starting from a weighted adjacency matrix *A* derived from STRING (24), we define the column-stochastic transition matrix Π_*ij*_ = *A*_*ij*_*/Σ* _*k*_ *A*_*kj*_ . For a protein index *g*, we define a restart vector *e*_*g*_ ∈ ℝ^*N*^, where (*e*_*g*_)_*i*_ = 1 if *i* = *g* and zero otherwise. The corresponding PPR vector is estimated iteratively as 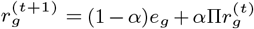 until convergence. The resulting vector *r*_*g*_ captures the steady-state visitation probabilities of random walks with restart from node *g*. Stacking these vectors across the interactome yields a high-dimensional positional representation. Because the raw PPR dimensionality is too large for direct model input, we reduce it with PCA before training, retaining *d*_PE_ principal components; the chosen *d*_PE_ for each model and benchmark is reported in Suppl. Tables 4 and 5. To fuse sequence and graph-derived features, we use a Feature-wise Linear Modulation (FiLM) layer (19). Two learned projections of the reduced positional encoding (PE) generate a feature-wise scaling term *γ*(PE) and shift term *β*(PE). Given the pooled sequence feature vector *x*, we apply a residual FiLM transformation *h* = *x* ⊙ (1 + *α γ*(PE)) + *α β*(PE), where ⊙ denotes element-wise multiplication and *α* is a learned scalar. The modulated representation is then passed to a classifier head to estimate the RNA-binding probability. This head is identical in the LoRA and FiLM settings: LayerNorm(*H*) → Dropout(0.2) → Linear(*H*, 2), where *H* is the pooled sequence-feature dimension. The linear layer outputs two logits (RBP vs non-RBP), converted to class probabilities with a softmax.

### Training Procedure

We train the model for binary classification using a class-weighted loss to account for label imbalance. In the LoRA setting, optimization updates the LoRA adapter parameters together with the task-specific FiLM and classification layers, while the frozen pLM backbone remains unchanged. Hyperparameter optimization was conducted via random search over the training hyperparameter space; the final settings used for each model and benchmark are summarised in Tables 4 and 5. To prevent overfitting and ensure an unbiased evaluation, we employ early stopping optimized for the Area Under the Precision–Recall Curve (AUPRC) on an internal validation set. Final performance metrics are subsequently reported exclusively on a strictly held-out test partition. To guarantee robustness across random initializations, we evaluate the model using repeated random sub-sampling validation (Monte Carlo cross-validation) over multiple independent seeds. For pairwise benchmark comparisons shown in Figure 2, statistical significance is assessed with paired two-sided t-tests on matched seeds/splits (i.e., paired runs sharing the same train/test partition). Benchmark comparisons against HydRA (12) are carried out on a curated difficult-positive subset of the HydRA test set—retaining only RBPs that lack any annotated RNA-binding domain—together with Precision at top *K* as an additional retrieval metric; the full evaluation protocol is described in Suppl. Section A.2.

**Figure 2.**
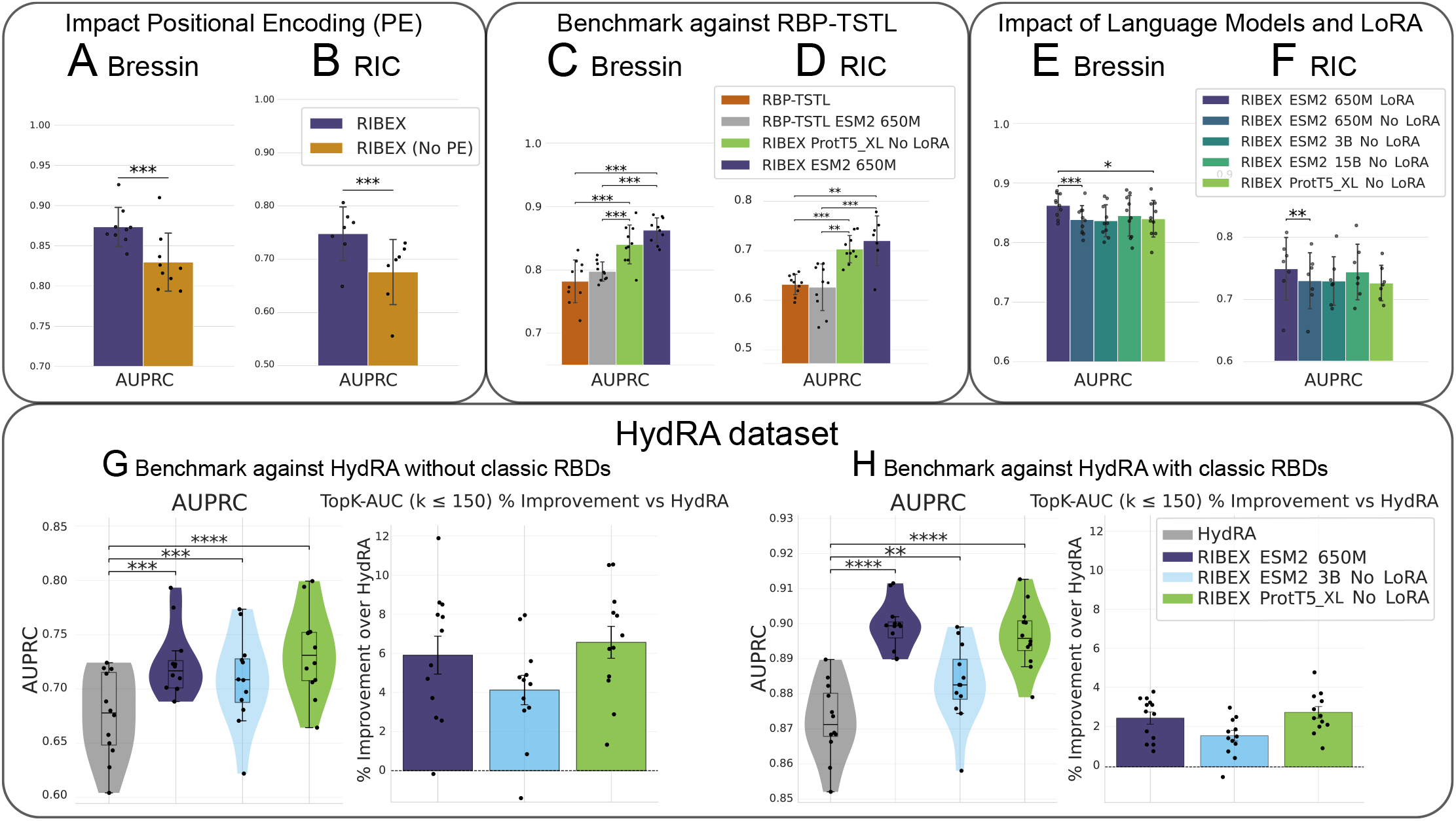
Benchmark Performance of RIBEX Across Datasets and Settings. (**A–B**) Effect of PPI-derived positional encodings (PEs) on the dataset by (5) (**A**) and the RIC dataset (**B**). (**C–D**) Comparison with RBP-TSTL on the dataset by (5) (**C**) and the RIC dataset (**D**). (**E–F**) Effect of pLM backbone choice and LoRA fine-tuning on the dataset by (5) (**E**) and the RIC dataset (**F**). (**G–H**) Comparison with HydRA on proteins without classical RNA-binding domains (**G**) and with classical RNA-binding domains (**H**); panels report AUPRC and the area under the Precision-at-top-*K* curve (Area Under the Precision-at-Top-*K* Curve (TopK-AUC), *k ≤* 150). Bars summarize runs/splits; asterisks denote paired two-sided *t*-test significance on matched seeds/splits (\**P <* 0.05, \*\**P <* 0.01, \*\*\**P <* 0.001).

### Model Interpretation

For the **sequence-level explanation via in silico Alanine scanning** (13; 1) of any given protein sequence, we first estimate its baseline RNA-binding probability, denoted *ŷ*. Then, we systematically substitute sliding windows of ten consecutive residues with alanine, advancing one residue at a time (stride 1). The fine-tuned language model is subsequently re-evaluated on each modified sequence, giving *ŷ*_A1a_. We measure the impact of the perturbation for each sequence window by the change in predicted probability Δ*ŷ*_A1a_ = *ŷ*_A1a_ − *ŷ*. A pronounced drop in the assigned probability (a highly negative Δ*ŷ*_A1a_) implies that the originally present residues are critical for conferring RNA-binding potential. Throughout this procedure, the positional encoding (PE) is held fixed at its original value, ensuring that any change in predicted probability stems from the sequence perturbation alone. Besides the sequence-level analysis, we investigate how the graph-derived positional encodings affect the model’s predictions (Figure 3). Keeping the amino acid sequence fixed, we perform **network-level feature ablation** by sequentially zeroing out individual dimensions of the reduced PE vector and measuring the resulting shift in the prediction score Δ*ŷ*_PE,*j*_ = *P*(*y* | *x*, **p**^(j)^) − *P*(*y*| *x*, **p**), where **p** ∈ ℝ^*d*^ denotes the original PE vector, **p**^(*j*)^ represents the same vector with its *j*-th dimension zeroed out, *x* is the input amino acid sequence, and *P* (*y* | ·) is the probability estimate from the trained classification network. PE dimensions that lead to a substantial decrease in the binding probability when disabled Δ*ŷ*_PE, *j*_ < − 0.01 are highlighted as carrying positive predictive evidence for RNA interaction. Because each reduced PE dimension is a PCA component, its effect is not directly interpretable at the level of individual interaction partners. We therefore collect, for each scanned protein, the PE dimensions whose ablation decreases the RNA-binding probability beyond the threshold above, set these dimensions to zero simultaneously, and apply the inverse PCA transform. This yields two reconstructed vectors in the full PPR-derived node space, one before and one after ablation, that is, a score over all proteins in the STRING interactome. We then compute the absolute node-wise difference between the original and ablated reconstructions and rank all nodes by this quantity. For each protein, we retain the top 40 nodes with the largest absolute differences and interpret them as the nodes most affected by the removed PE signal. Across proteins, we aggregate how often each node appears in these per-protein top-40 lists, which produces a frequency-based map of interactome regions that recurrently support positive predictions. For visualization, these aggregated node frequencies are overlaid on a t-distributed Stochastic Neighbor Embedding (t-SNE) embedding of the reduced PE matrix for all STRING proteins (Figure 3) and Suppl. Figure 4.

**Figure 3.**
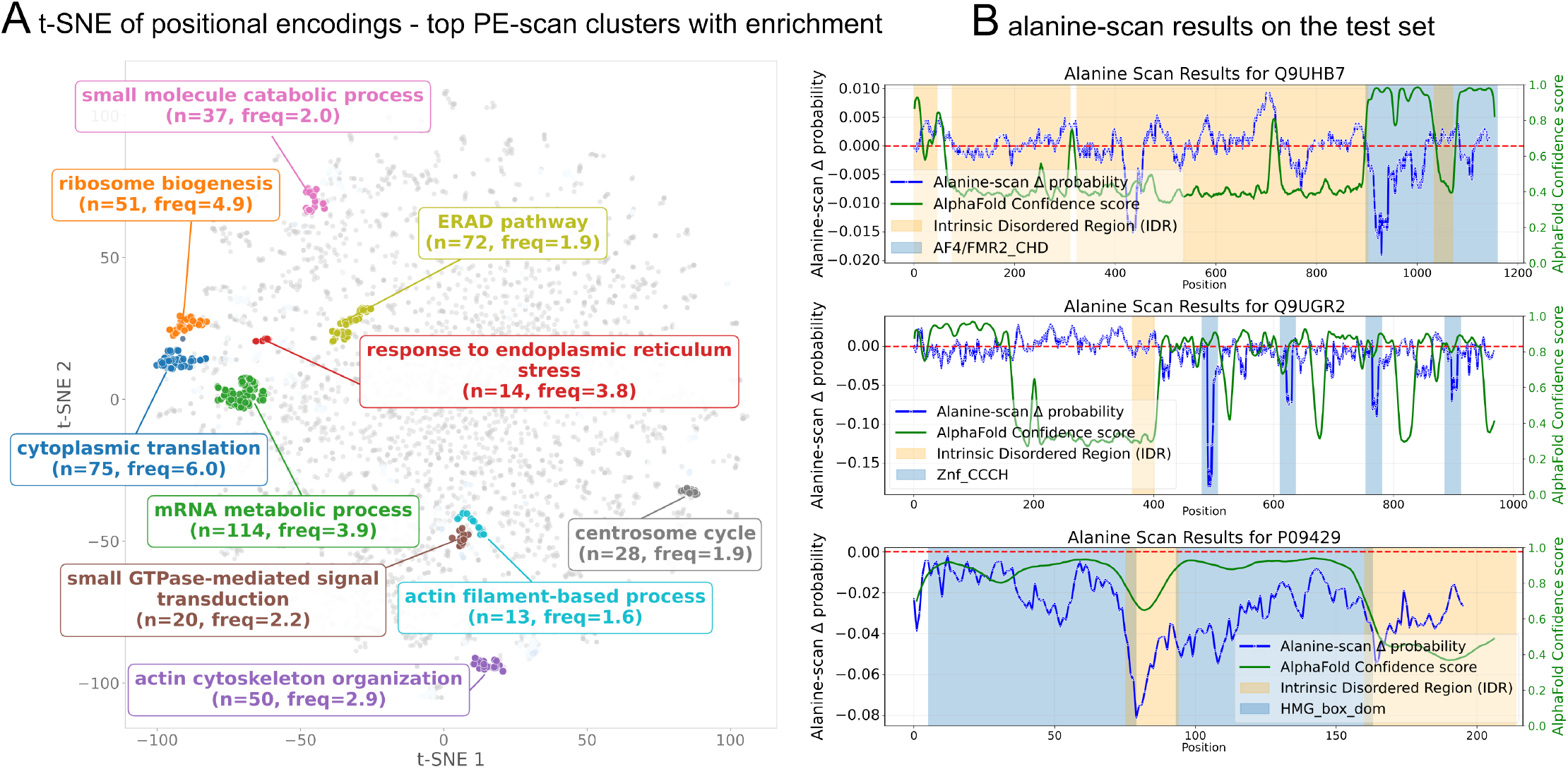
Explainability Analysis of RIBEX Predictions. **(A)** DBSCAN clustering of influential nodes in the t-SNE space. Colors indicate clusters; the legend reports cluster size (*n*), mean perturbation frequency (freq), and the top enriched Gene Ontology (GO) biological-process term. **(B)** Sequence-level computational alanine scans for three representative correctly predicted RBPs. The blue trace shows the per-window probability change Δ*ŷ*_A1a_, the red line marks the zero baseline, and the green trace shows smoothed AlphaFold pLDDT. Colored tracks indicate InterPro domain annotations and MobiDB IDRs.

## Results

### Impact of design choices

To disentangle the **contribution of the pLM** from that of parameter-efficient adaptation, we compared RIBEX with LoRA on ESM2-650M against RIBEX without LoRA using ESM2-650M, ESM2-3B, ESM2-15B, and ProtT5-XL backbones (Figure 2E for the dataset by (5); Figure 2F for the RIC dataset). Across both datasets, adding LoRA to ESM2-650M significantly improves performance over the untuned model, with the largest gains in Matthews Correlation Coefficient (MCC) (Suppl. Figure 6E,F) and AUPRC. In contrast, varying the pretrained backbone and scaling model size without LoRA yields only modest differences: a slight trend appears on the dataset from (5) but is weak and does not transfer clearly to the more challenging RIC benchmark. All in all, these results indicate that **parameter-efficient adaptation contributes more strongly to performance than increasing backbone size alone**. This pattern is consistent with recent analyses showing that, although pretrained protein language models outperform naive sequence representations, performance on many downstream tasks does not reliably scale with pretraining/model size and often depends on low-level features learned early in pretraining (14).

To assess the **contribution of positional encodings (PEs)**, we compare RIBEX with and without PEs in two benchmark settings. In panel A of Figure 2, corresponding to the dataset by (5), the PE-enabled model consistently outperforms the no-PE variant. The same trend is observed in panel B on the human RIC dataset, where adding PEs to the LoRA-fine-tuned human model yields higher scores across the reported evaluation metrics. This indicates that **positional information provides a measurable benefit**.

### Benchmarking Against State-of-the-Art Methods

We benchmark RIBEX against two classes of state-of-the-art RBP prediction methods. First, **RBP-TSTL** (18), which to our knowledge represents the current state of the art in RBP prediction using pLM representations, also shown to outperform earlier non–deep learning approaches such as TriPepSVM. Second, we compare against network-based and hybrid approaches, including **SONAR 3.0** (4), which predicts RBPs by exploiting the observation that proteins interacting with multiple RBPs are themselves likely to bind RNA, and HydRA (12), a hybrid deep-learning framework that integrates protein sequence features with PPI network information.

RIBEX with ProtT5-XL embeddings and PEs already significantly outperforms the baseline RBP-TSTL configuration even without LoRA fine-tuning, showing relative AUPRC improvements of 7.5% and 10.8% on the dataset from (5) and RIC benchmarks, respectively (Figure 2C,D), and corresponding relative MCC gains of 7.1% and 9.3% (Suppl. Figure 6C,D). Replacing ProtT5-XL with ESM2-650M slightly improves the RBP-TSTL baseline, particularly on the dataset from (5). However, the RIBEX architecture consistently outperforms both configurations across all benchmarks, with significant AUPRC improvements (Figure 2) and corresponding MCC gains (Suppl. Figure 6C,D). Motivated by the RBP-TSTL design, we also tested multi-species pretraining in our pipeline using *E. coli, Salmonella, Mus musculus*, and *Saccharomyces cerevisiae*; however, this strategy did not improve performance in our setting (see Suppl. Section B.2).

**On the HydRA benchmark** (12), we run RIBEX on the same dataset and compare against the published test set predictions distributed with HydRA, including HydRA-seq (the sequence-only variant with ablated network information) and SONAR 3.0 (4). Details of the evaluation protocol and metrics are provided in Suppl. Section A.2, with additional plots in Suppl. Section B.3. Panels G and H of Figure 2 report AUPRC together with TopK-AUC, defined as the area under the Precision-at-top-*K* curve for *k* ≤ 150. In this setting, both RIBEX variants—ESM2-650M with LoRA and ProtT5-XL without LoRA—outperform the HydRA baseline. For AUPRC, relative improvements over HydRA are approximately 5.9% and 6.6% on the subset lacking classical RBDs for ESM2-650M and ProtT5-XL, respectively, and approximately 2.5% and 2.7% on the classical-RBD subset (Figure 2G,H). Notably, for proteins lacking classic RNA-binding domains (panel G) the relative drop in performance is smaller for RIBEX than for HydRA when compared with the classic-RBD subset (panel H), indicating improved robustness on this harder subset.

### Explainability

To probe which aspects of the model’s learned representation drive its predictions, we apply two complementary perturbation-based analyses detailed in Section 2.5: a network-level positional encoding scan (Figure 3A) and a sequence-level computational alanine scan (Figure 3B).

#### Positional Encoding Scan Identifies Functional Interactome Communities

Applying the network-level feature ablation strategy (Section 2.5) to high-confidence true-positive RBPs from the test set yielded a per-node frequency score. When these scores are overlaid on a t-SNE embedding of the PE matrix (Figure 3A), the most frequently appearing neighbour nodes form spatially coherent clusters rather than being diffusely distributed across the projection. This suggests that the proteins most frequently affected by the PE perturbation analysis are not randomly distributed in the learned network-context space, but instead seem to concentrate in regions associated with coherent functions. To delineate these regions, we applied DBSCAN, a density-based clustering method, to the t-SNE coordinates of nodes with non-zero perturbation frequency, followed by GO biological-process enrichment on each cluster (Figure 3A). Enriched cluster annotations include processes where RBPs play key roles, such as cytoplasmic translation and ribosome biogenesis, as well as cytoskeleton organization, ER stress–related pathways involving localized translation, and centrosome and cell-cycle functions. Together, these observations support the interpretation that RIBEX leverages structured interactome communities rather than isolated graph features when assigning RNA-binding propensity.

#### Alanine Scanning Resolves Domain and IDR Contributions

To complement the network-level analysis, we applied *in silico* alanine scanning (Section 2.5) to a small set of correctly predicted RIC-positive test proteins, prioritizing proteins with extensive IDR content. The resulting per-window probability changes (Δ*ŷ*_A1a_), where strong negative peaks indicate important sequence segments, are visualized together with smoothed AlphaFold confidence scores and domain/IDR annotations (Figure 3B). Importantly, this analysis operates on full-length sequences, capturing perturbation effects in both ordered domains and disordered regions without requiring prior structural information.

As an example, we highlight **Q9UHB7**, the ALF transcription elongation factor 4 (AFF4), an intrinsically disordered scaffold protein of the Super Elongation Complex (SEC) that has also been reported to bind RNA. In addition to extensive IDRs, AFF4 contains a conserved C-terminal homology domain (C-terminal Homology Domain (CHD)) that mediates AFF-family dimerization and binds RNA and DNA *in vitro* (7). Consistent with this function, the strongest perturbation signal in Figure 3B localizes to the C-terminal CHD, with additional decreases in disordered regions. Although these sites mediate protein–protein interactions rather than direct RNA contacts, they may contribute to RNA-associated predictions through complex context.

The alanine scan of **Q9UGR2**, a regulator of miRNA biogenesis, shows a sharply localized perturbation overlapping the annotated CCCH-type zinc-finger RNA-binding domain (ZnF CCCH), while flanking regions display minimal effects, indicating that the model concentrates its sequence-level evidence on this compact RNA-binding module.

In **P09429 (HMGB1)**, a multifunctional chromatin-associated protein with RNA-binding activity, the alanine scan reveals the largest drop in predictive probability at the IDR boundary between the A-box and the adjacent hinge (∼70–90), with additional sensitivity in the unstructured acidic tail (aa 186–215). Although **HMGB1** is known to bind structured RNAs (22), precise interaction sites remain unclear. Studies suggest this disordered segment may act as a structural hinge positioning the two HMG boxes for coordinated RNA binding rather than contacting RNA directly (22). Additional examples are provided in Suppl. Figure 9 and section C.2, including similar HMGB-family profiles for P26583 (HMGB2) and O15347 (HMGB3) and a complementary BUD13 (Q9BRD0) case in which an alanine-sensitive IDR segment is consistent with a RES-complex partner interface rather than a direct RNA-binding motif.

Together, these *in silico* profiles illustrate how RIBEX uses both classical domains and essential IDR linkers to evaluate RNA-binding propensity.

## Discussion

Here we present RIBEX, a novel method for predicting both canonical and non-canonical RBPs by integrating protein sequence information with network topology. Across benchmarks, positional encodings (PE) consistently improve performance, supporting our central premise that network topology provides information complementary to sequence features. Accordingly, RIBEX remains competitive with or outperforms strong baselines such as RBP-TSTL and HydRA (Figure 2C,D,G,H), despite using a comparatively simple FiLM-based fusion mechanism.

**Ablation results** show that task-specific LoRA adaptation on ESM2-650M improves performance more than simply scaling the frozen pLM backbone (Figure 2E,F). The limited effect of backbone size, especially on the challenging RIC benchmark, indicates that human-only RBP prediction depends not just on sequence representation but also on task adaptation and contextual information, consistent with the additional gains from PE after LoRA fine-tuning. **Dataset differences** are informative: (5) is annotation-based, enriched for canonical RBPs, while the RIC dataset captures experimentally detected RNA associations, including indirect or context-dependent interactions. The stronger impact of topology and improved robustness on non-canonical RBPs in the HydRA benchmark is thus biologically plausible, as interactome context helps reveal RNA-associated proteins beyond sequence signals. The **interpretability analyses** further support this view. At the sequence level, computational alanine scanning highlights sensitivity in known RNA-binding domains, as well as selected IDR segments (Figure 3B), while network-level PE ablation reveals recurrent, functionally coherent interactome communities linked to positive predictions (Figure 3A). Importantly, these analyses support the conclusion that the model leverages broader RNA-associated cellular context, and not necessarily direct RNA binding, as shown for the AFF4 example.

Limitations of RIBEX include its focus on protein-level RNA-binding predictions without resolving RNA target specificity or residue-level contacts, its evaluation being restricted to human proteins and a single PPI graph (STRING), and the network explanation pipeline providing high-level summaries of influential neighborhoods rather than causal explanations. In addition, sequence-level attributions from the alanine scan map model sensitivity along the protein, with peaks highlighting domains or IDR/interface regions that contribute to predicted RNA association which are not necessarily direct.

Future improvements could include tighter sequence–graph integration via pLM-based node features or cross-modal message passing, alternative PEs, and extensions to cross-species or protein–RNA interaction prediction, all aimed at boosting predictive accuracy and supporting more interpretable, subgraph-level reasoning. Taken together, our results position RIBEX as a practical framework for prioritizing candidate RBPs and generating mechanistic hypotheses, with complementary interpretability particularly valuable for non-canonical proteins where sequence alone is insufficient.

## Data and Code Availability

The code and documentation necessary to reproduce our results, alongside links to the used datasets, are available at https://github.com/marsico-lab/RIBEX.

## Author contributions statement

S.F. and F.S. conceived the study and designed the methodology. F.S. developed the initial implementation and performed dataset curation and preprocessing. S.F. designed and conducted the experiments and analyses. S.F., F.S. and A.M. wrote the manuscript. G.K. and A.M. provided funding. A.M. and M.H. did the primary supervision. G.K. and M.H. revised the manuscript. All authors reviewed and approved the final manuscript.

## Acknowledgments

S.F. were supported by the Helmholtz Association under the joint research school Munich School for Data Science—MUDS; F.S and G.K are funded by the Technical University of Munich. Further support to A.M. was obtained by the SFB TRR267 (ID: 403584255, project Z02) and the Deutsche Forschungsgemeinschaft (DFG, German Research Foundation) – Project-ID 533767322 – EXC 3113/1, Cluster for Nucleic Acid Sciences and Technologies – NUCLEATE.

