## Supplementary for "RIBEX: Predicting and Explaining RNA Binding Across Structured and Intrinsically Disordered Regions (IDR)-rich Proteins"

#### Methods and Experimental Details

##### Dataset Split Statistics

The tables below report the size and class composition of the training, validation, and test partitions used for each benchmark. Values are representative of a single seed (seed 12345); partition sizes are identical across all ten seeds by construction, since the common gene pool is fixed. Positive counts refer to proteins labelled as RBPs.

**Table 1 Split statistics for the RIC benchmark.** The common gene pool consists of 15,299 proteins present in the embeddings of all four evaluated pLMs. The hold-out set (10%) is divided into a validation fold (one third, used for epoch selection) and a test fold (two thirds, used for final evaluation).

| Split | Samples | Positives | Positive fraction |
|---|---|---|---|
| Train | 13,769 | 1,430 | 10.4% |
| Validation | 510 | 48 | 9.4% |
| Test | 1,020 | 96 | 9.4% |
| *Total* | *15,299* | *1,574* | *10.3%* |

**Table 2 Split statistics for the dataset from (2).** The common gene pool consists of 10,414 proteins present in the embeddings of all four evaluated pLMs. The hold-out set (10%) is divided into a validation fold (one third) and a test fold (two thirds).

| Split | Samples | Positives | Positive fraction |
|---|---|---|---|
| Train | 9,372 | 1,208 | 12.9% |
| Validation | 347 | 47 | 13.5% |
| Test | 695 | 93 | 13.4% |
| *Total* | *10,414* | *1,348* | *12.9%* |

**Table 3 Split statistics for the HydRA benchmark, stratified by pLM.** Training data comprises all proteins in the HydRA repository dataset that are not part of the official test set. The test set is taken directly from the HydRA repository; sizes vary across pLMs because only proteins for which embeddings are available are retained for training, while for the test set the size is the same, corresponding to the test samples available across all models. Per-seed validation proteins for epoch selection are derived by bootstrap resampling with replacement from the test set (out-of-bag proteins) and are not listed as a separate partition.

| Model | Train | Train positives | Test | Test positives |
|---|---|---|---|---|
| ESM2 650M | 12,645 | 2,459 | 2,994 | 575 |
| ESM2 3B | 12,381 | 2,358 | 2,994 | 575 |
| ProtT5-XL | 12,974 | 2,588 | 2,994 | 575 |

##### Benchmark Evaluation Protocol

All comparisons against HydRA (5), HydRA-seqOnly, and SONAR 3.0 (1) are performed on the HydRA held-out test set. For each independent random seed, we restrict all models to their common protein coverage by computing the intersection of proteins available in every compared model. This shared set is then split into a validation subset (30%) and a test subset (70%) using stratified random sampling to preserve the positive class fraction. For each RIBEX variant, the training checkpoint that maximises AUPRC on the validation subset is selected; predictions from this checkpoint are then generated on the held-out test subset. Baseline scores for HydRA, HydRA-seqOnly, and SONAR 3.0 are fixed post-hoc predictions and are evaluated directly on the same test subset.

###### Difficult-positive filter.

Canonical RNA-binding domains (RBDs) such as RRM or KH domains are strong and easily detectable sequence signals; proteins carrying them are straightforwardly classified as RBPs by almost any sequence-based method. To probe each model's capacity to detect RNA binding in the absence of such canonical features—which is particularly relevant for IDR-mediated RNA association—we apply a *difficult-positive* filter to the test set. Positive proteins are retained only if they are annotated as RBPs in the curated catalogue (3) *and* carry no annotated RBD (i.e., zero known RNA-binding domains, zero RBD sequence coverage). All negative proteins are kept without modification. Easy positives — those whose RNA-binding activity is straightforwardly predicted from canonical domain content — are thereby excluded from the evaluation.

###### Evaluation metrics.

Two retrieval metrics are computed on the filtered test set. The **area under the precision-recall curve (AUPRC)** summarises overall discriminative performance under class imbalance; because positives are a small minority of the human proteome, AUPRC is more informative than Area Under the Receiver Operating Characteristic Curve (AUROC) in this setting. **Precision at top** $K$ (Precision at Top $K$ (P@K)) measures the fraction of true RBPs among the $K$ highest-scoring proteins for $K \in [10, 300]$, directly quantifying the practical utility of each model for prioritising experimental follow-up. The area under the P@K curve (TopK-AUC) is computed by numerical integration of P@K over

$K$ after linearly rescaling $K$ to $[0, 1]$, so that the score summarizes ranking quality across the selected cutoff range. In the main-text HydRA comparison (Figure 2G,H), we report the restricted variant computed over $K \leq 150$ to emphasize the regime most relevant for actionable candidate lists.

##### Statistical testing.

Comparisons are performed over twelve independent random seeds. Seed-matched metric pairs (i.e., values from the same partition) are compared with a two-sided paired $t$-test. When multiple models are tested against HydRA simultaneously, $p$-values are adjusted with the Holm–Bonferroni procedure (4).

#### Hyperparameters

Tables 4 and 5 report the final hyperparameters used for all RIBEX model variants. For the RIC and (2) datasets, hyperparameters were held fixed; for the HydRA benchmark, LoRA hyperparameters were determined by random search and are model-specific. FiLM PE settings are identical across all four pLMs within each benchmark. All LoRA adapters target the key and value projection matrices of the self-attention layers.

**Table 4 FiLM PE hyperparameters.** Settings are shared across all pLMs (ESM2 650M, ESM2 3B, ESM2 15B, ProtT5-XL) within each benchmark. $d_{\mathrm{PE}}$: number of PCA components retained for the positional encoding. Patience refers to the early-stopping patience in epochs (monitored on validation AUPRC).

| Benchmark | Learning rate | Batch size | $d_{\mathrm{PE}}$ |
|---|---|---|---|
| RIC / dataset from (2) | $5.0 \times 10^{-4}$ | 256 | 512 |
| HydRA | $9.3 \times 10^{-5}$ | 256 | 512 |

**Table 5 LoRA hyperparameters.** For RIC and the dataset from (2), the same settings are used for both ESM2 650M and ProtT5-XL; rank, $\alpha$, and dropout were left at default values in those benchmarks. For HydRA, hyperparameters were optimised separately per backbone via random search. $r$: LoRA rank; $\alpha$: LoRA scaling factor; WD: weight decay; $d_{\mathrm{PE}}$: PCA components for positional encoding. Patience refers to early-stopping patience (validation AUPRC).

| Benchmark | Model | LR | BS | $d_{\mathrm{PE}}$ | $r$ | $\alpha$ | Dropout | Weight Decay |
|---|---|---|---|---|---|---|---|---|
| RIC / dataset from (2) | ESM2 650M | $3.0 \times 10^{-4}$ | 256 | 512 | 3 | 0.42 | 0.45 | $5.3 \times 10^{-4}$ |
| | ProtT5-XL | $3.0 \times 10^{-4}$ | 256 | 512 | 3 | 0.42 | 0.45 | $5.3 \times 10^{-4}$ |
| HydRA | ESM2 650M | $9.1 \times 10^{-4}$ | 415 | 418 | 6 | 1.66 | 0.38 | $3.7 \times 10^{-4}$ |
| | ProtT5-XL | $3.4 \times 10^{-4}$ | 392 | 466 | 10 | 1.58 | 0.38 | $1.6 \times 10^{-4}$ |

#### Positional Encoding Experiments

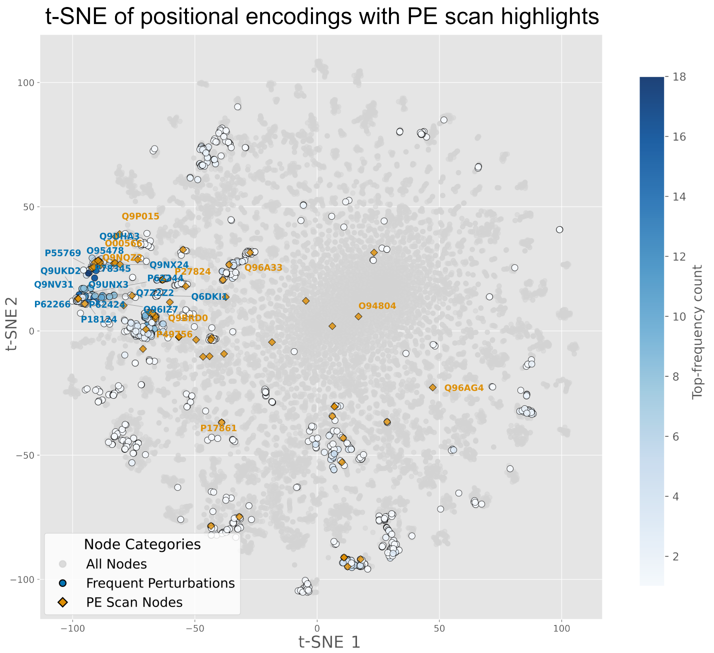


**Figure 4 Frequency map of the positional-encoding explanation analysis.** Nodes are shown in the reduced PE t-SNE space together with the aggregated frequency counts from the inverse-PCA PE-scan analysis. Blue points denote interacting partners, and orange points denote the top predicted positives in the held-out test set.

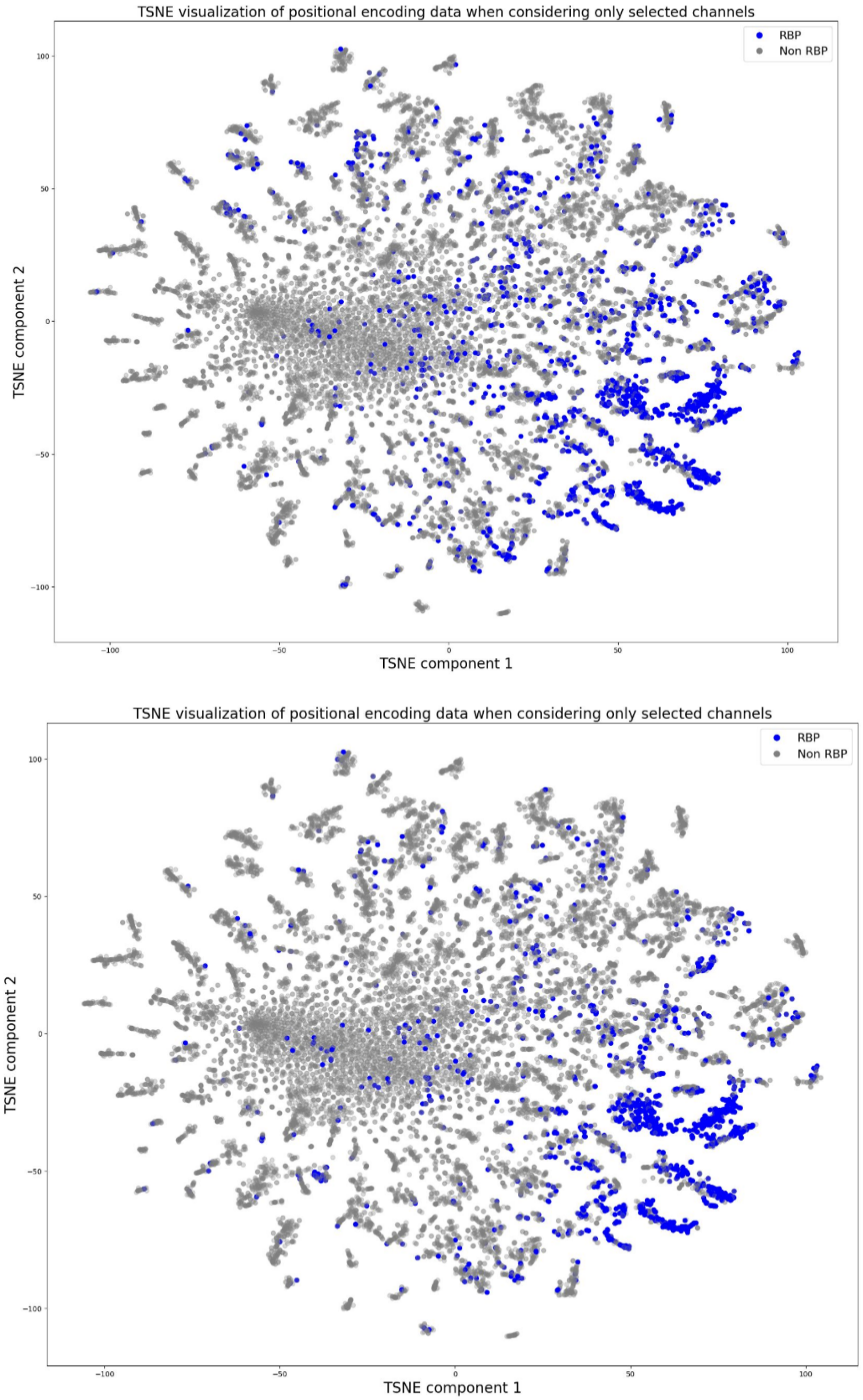


**Figure 5 t-SNE Visualization of Positional Encoding Data for Selected Channels. Top:** dataset from (2) positional-encoding subset. **Bottom:** RIC positional-encoding subset.

#### Additional Results

##### Additional Benchmark Figures

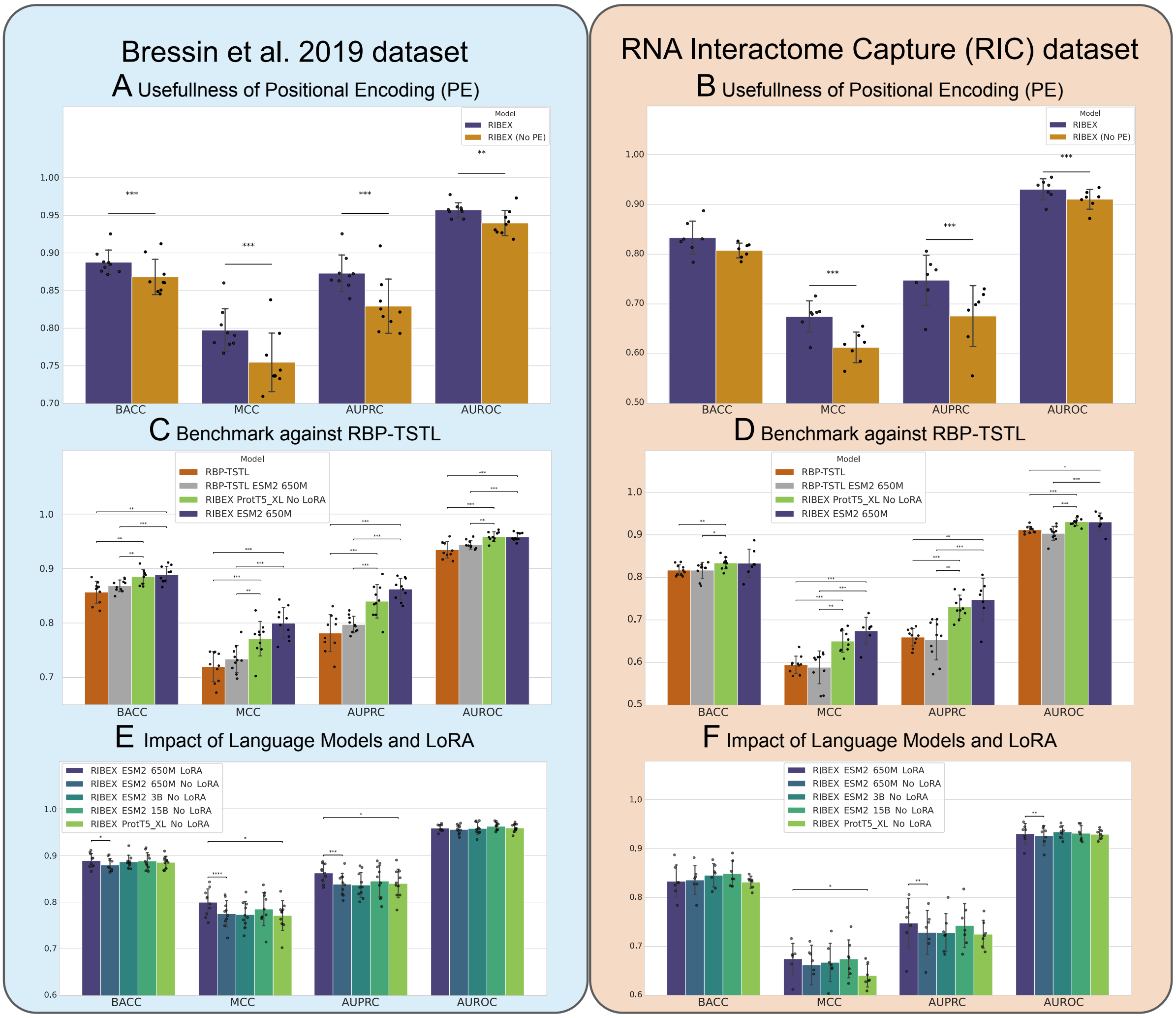


**Figure 6 Full benchmark comparison across all reported metrics.** Panels **A–B** show the effect of positional encodings on the dataset from (2) and RIC benchmarks, respectively. Panels **C–D** compare RIBEX against RBP-TSTL on the dataset from (2) and RIC. Panels **E–F** show the effect of pLM backbone choice and LoRA fine-tuning on the dataset from (2) and RIC. This expanded supplementary figure includes the full metric set used during benchmarking and is therefore referenced in the main text for metrics not shown in Figure 2.

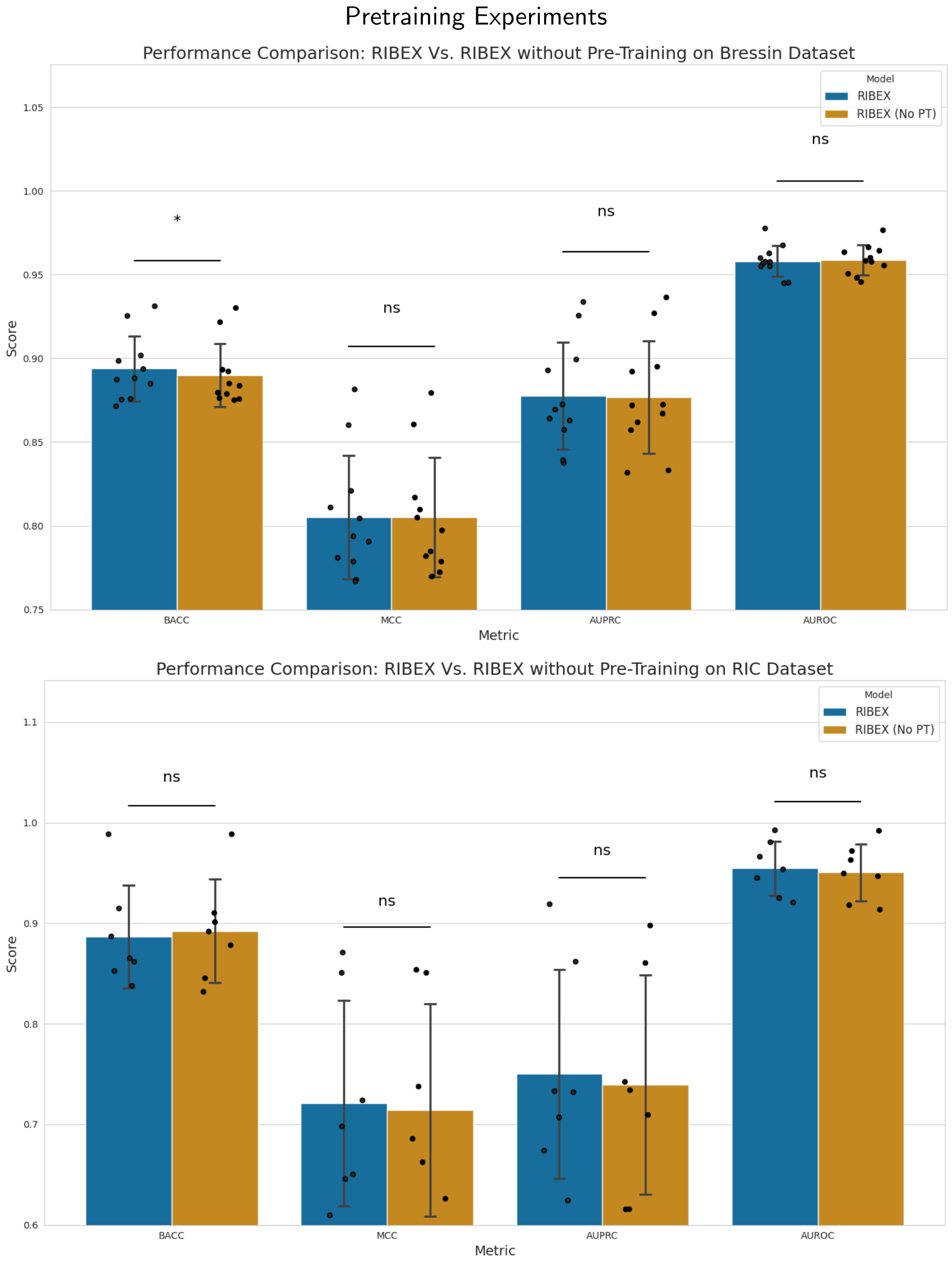


**Figure 7 Performance Comparison Between RIBEX and RIBEX Without Pre-Training. Top:** dataset by (2). **Bottom:** RIC dataset

#### HydRA Supplementary Figures

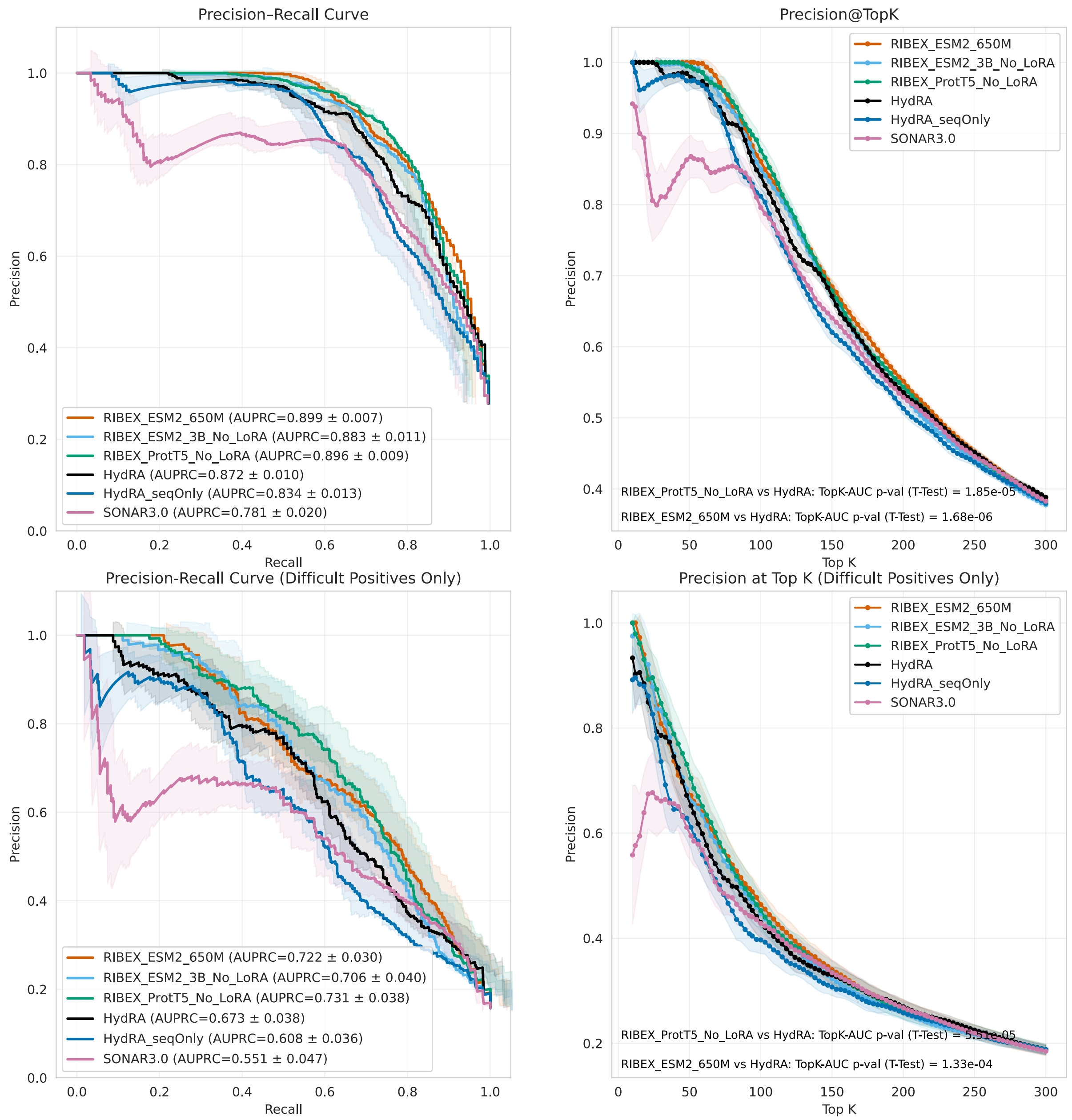


**Figure 8 AUPRC and Precision-at-Top-$K$ Comparison Between HydRA (5), HydRA-Sequence-Only, SONAR 3.0 (1), and RIBEX.** Using a paired $t$-test to compare against RIBEX (ESM2, 650M), RIBEX (ESM2, 3B) without LoRA, and RIBEX (ProtT5) without LoRA. Comparison is done on all test-set positives (**Top**) and on only those not showing any canonical RNA-binding domain (**Bottom**).

### Interpretability Analyses

#### Additional Alanine Scan Results

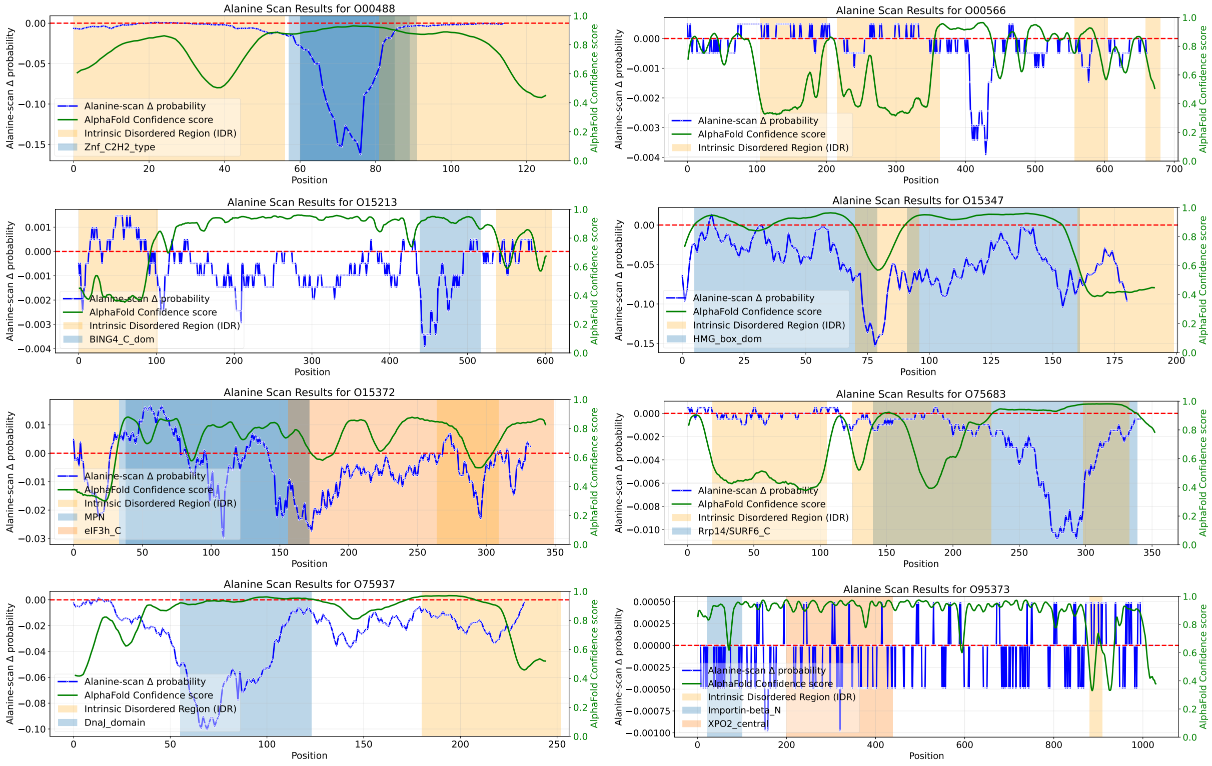


**Figure 9 Alanine-scan attribution maps for selected proteins.** Each panel shows the alanine-scan–based importance profile for the corresponding UniProt ID. The figure continues on the following pages.

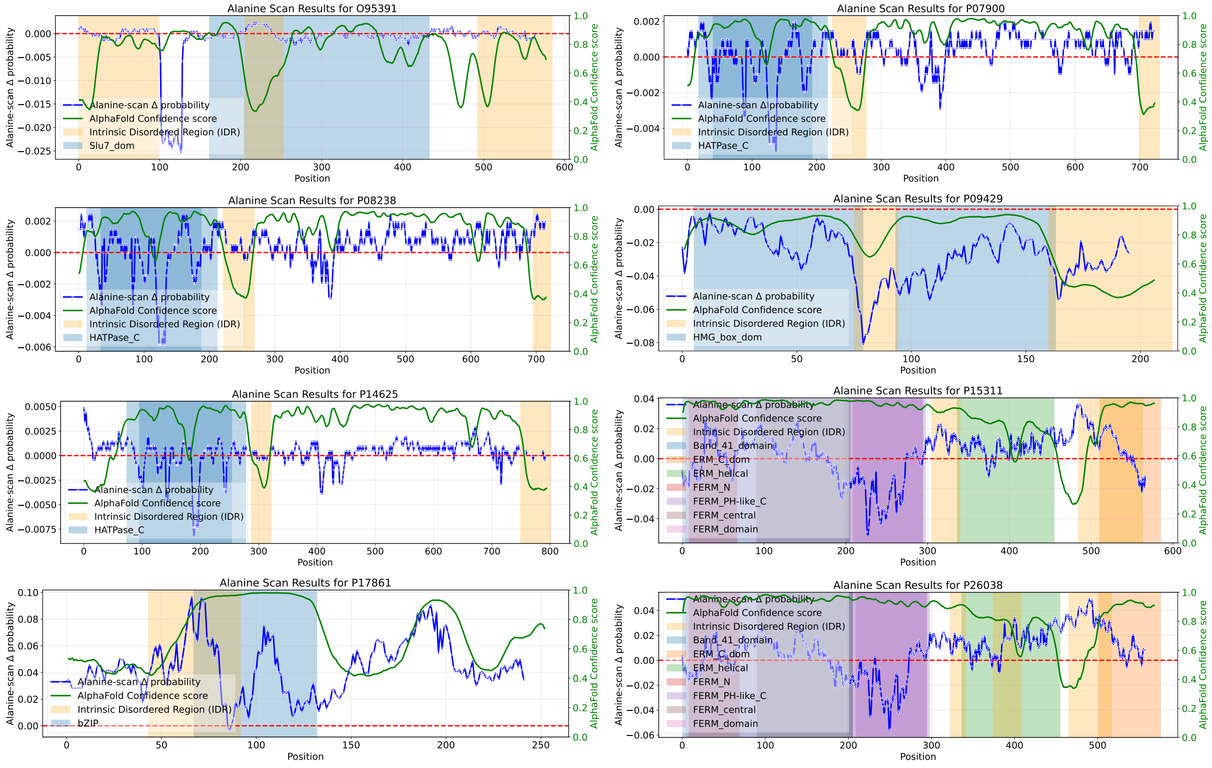


**Figure 10 Alanine-scan attribution maps for selected proteins (continued).**

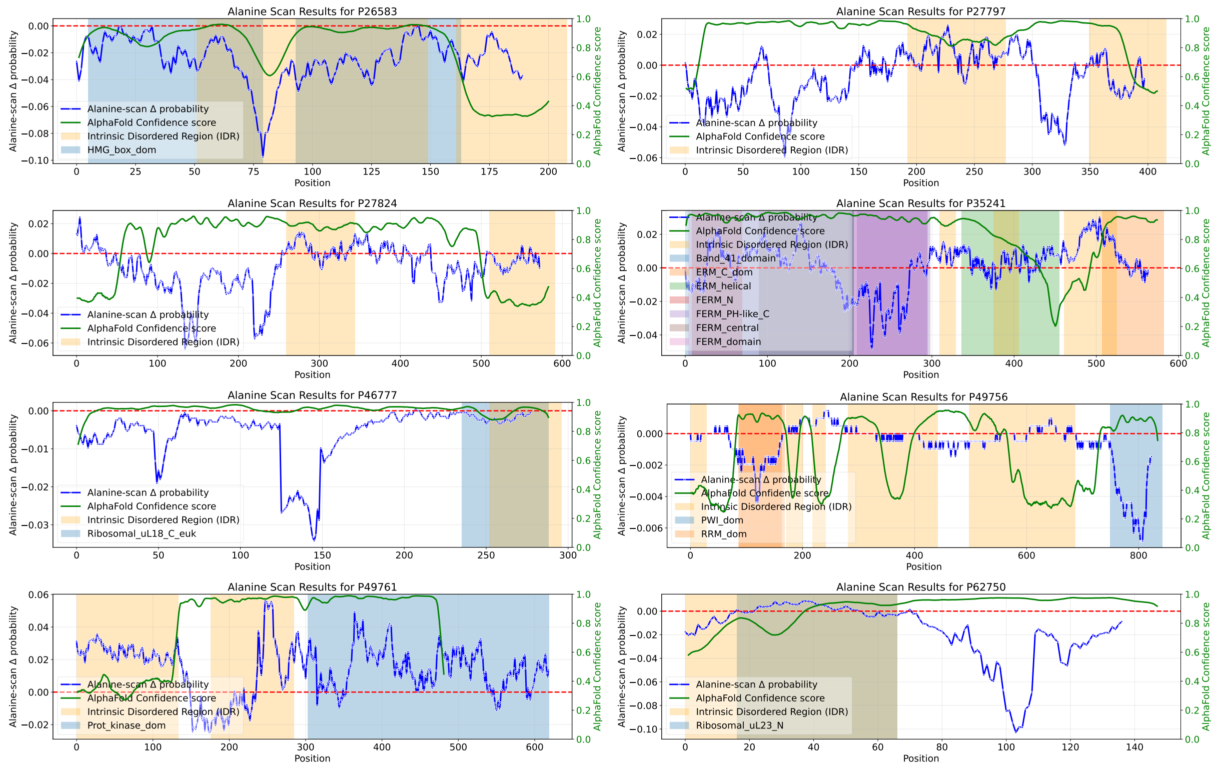


**Figure 11 Alanine-scan attribution maps for selected proteins (continued).**

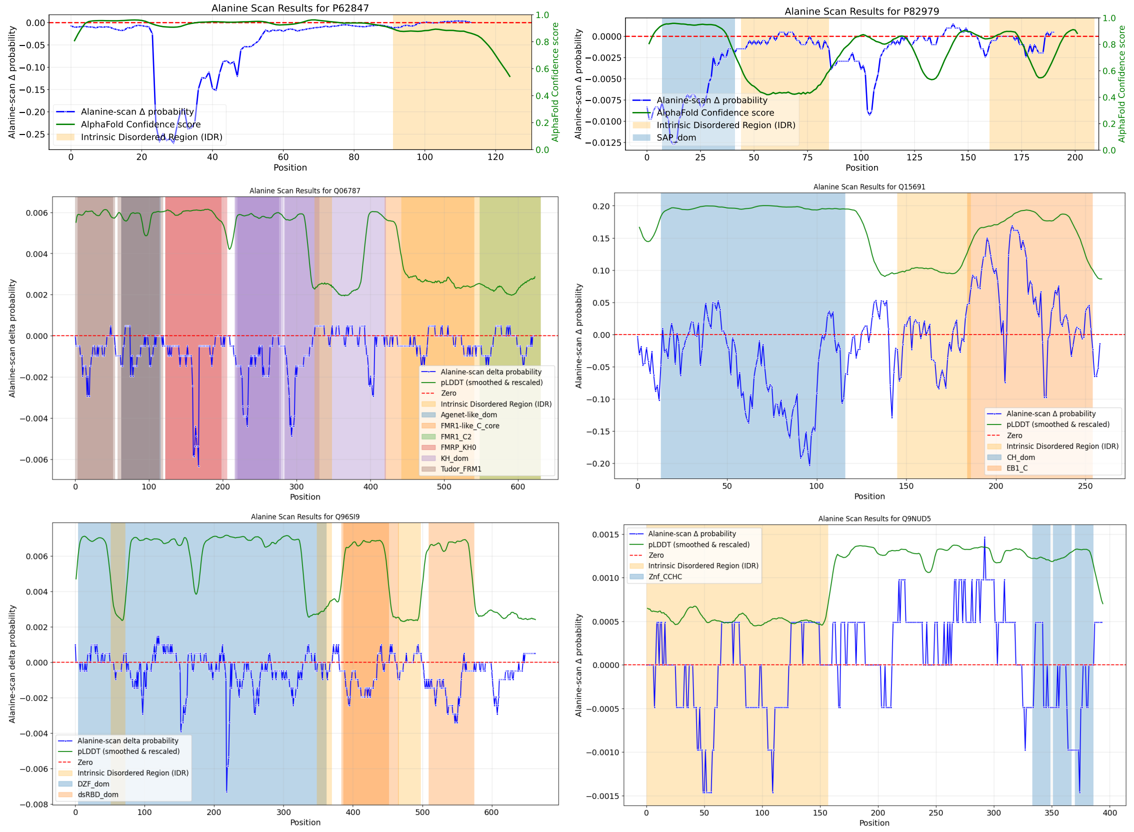


**Figure 12 Alanine-scan attribution maps for selected proteins (continued).**

##### Additional HMGB-family examples.

Beyond HMGB1 in the main text, the attribution maps in Figures 9 and 13 show qualitatively similar profiles for P26583 (HMGB2) and O15347 (HMGB3). In both cases, the signal extends across the flexible linker/IDR context and into the acidic C-terminal tail, consistent with a broader contribution of disordered inter-domain regions to the model's RNA-associated predictions. These related profiles support the interpretation that, across HMGB-family proteins, such segments can shape the prediction without necessarily corresponding to direct RNA-contacting motifs.

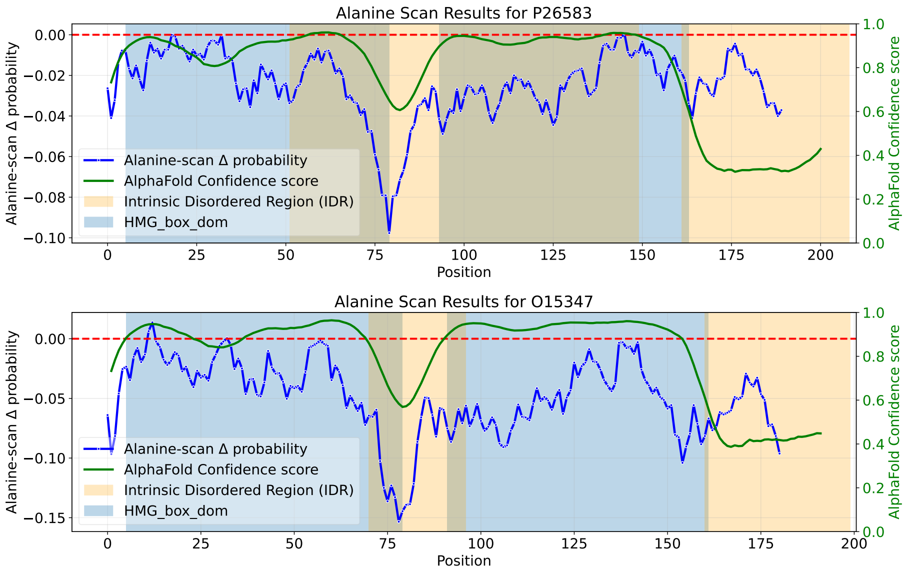


**Figure 13 Focused HMGB-family alanine-scan attribution maps.** The attribution profiles for P26583 (HMGB2; left) and O15347 (HMGB3; right) highlight signal spanning flexible linker/IDR context and the acidic C-terminal region, complementing the HMGB1 example discussed in the main text.

#### Supplementary Structural Hypothesis for BUD13 (Q9BRD0)

The main text cites a localized alanine-scan sensitivity in human BUD13 (Q9BRD0) around residues ∼580–590. Here we show the corresponding attribution profile and provide a structural interpretation as a hypothesis, not a residue-level assignment.

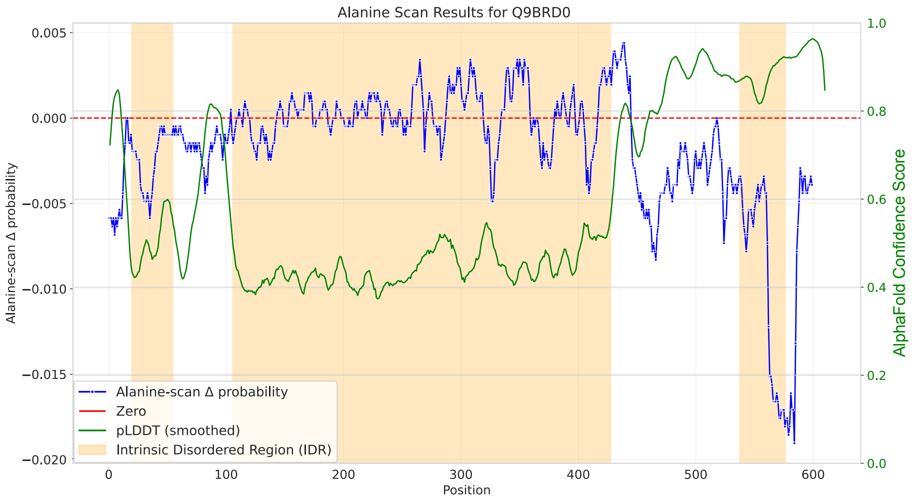


**Figure 14 BUD13 (Q9BRD0) alanine-scan attribution profile.** The strongest signal localizes around residues ∼580–590 in an annotated IDR, consistent with a candidate partner-binding segment in the RES-complex context.

By analogy to the yeast Snu17p–Bud13p RES-subcomplex structure (PDB 4UQT), in which a short Bud13p peptide binds the Snu17p RRM/UHM-like surface through a compact protein–protein interface (6), the human BUD13 signal is compatible with a partner-binding segment involved in RES-complex assembly. This is also consistent with the signal falling in an annotated IDR with locally elevated pLDDT relative to the surrounding sequence. As an additional motif-level observation, an `RWDGV` sequence is present in human BUD13 near residues 583–587, matching a motif reported for the yeast ligand. This correspondence is suggestive, but it does not establish residue-level equivalence or conserved binding geometry.
